# Glucose regulates expression of pro-inflammatory genes *IL-1β* and *IL-12* through a mechanism involving hexosamine biosynthesis pathway dependent regulation of α-E catenin in the RAW 264.7 macrophage cell line

**DOI:** 10.1101/2021.04.14.439728

**Authors:** Waruni C. Dissanayake, Jin Kyo Oh, Brie Sorrenson, Peter R. Shepherd

## Abstract

High glucose levels are associated with changes in macrophage polarization and evidence indicates that the sustained or even short-term high glucose levels modulate inflammatory responses in macrophages. However, the mechanism by which macrophages can sense the changes in glucose levels are not clearly understood. We find that high glucose levels rapidly increase the α-E catenin protein level in RAW264.7 macrophages. We also find an attenuation of glucose induced increase of α-E catenin when hexosamine biosynthesis pathway is inhibited either with glutamine depletion or with the drugs azaserine and tunicamycin. This indicates the involvement of hexosamine biosynthesis pathway in this process. Then, we investigated the potential role of α-E catenin in glucose induced macrophage polarization. We find that the reduction of α-E catenin level using siRNA attenuates the glucose induced change of IL-1β mRNA level under LPS stimulated condition. Further, we identified that the depletion of α-E catenin also decreases the *IL-12* gene expression in basal glucose conditions leading to a reduction of glucose induced changes in *IL-12*. Together this indicates that α-E catenin can sense the changes in glucose levels in macrophages *via* hexosamine biosynthesis pathway and also can modulate the glucose induced gene expression of inflammatory markers such as *IL-1-β* and *IL-12*. This identifies a new part of the mechanism by which macrophages are able to respond to changes in glucose levels.

## Introduction

Macrophages are highly sensitive to environmental stimuli. To acquire distinct functional phenotype against host cell infection, macrophages can be polarized into either pro-inflammatory cytokine secreting-classically activated (M1) or anti-inflammatory cytokine secreting-alternatively activated (M2) phenotype. A sustained or even short-term exposure to high glucose levels is known to induce M1 macrophage polarization [1] [2]. This has potential to be relevant in disease such where derangements in glucoregulatory mechanisms play an important role. One such disease is COVID-19 as elevated glucose levels favour SARS-CoV2 infection and effects of glucose levels on macrophage function have been shown to play an important role in this [3, 4]. Another example is type-2 diabetes where hyperglycaemia leads to the accumulation of macrophages and other innate immune cells with M1 phenotype [5]. Moreover, it has been shown that M1 macrophages highly depend on glycolysis as the energy source [6, 7], while M2 macrophages rely on fatty acid oxidation [8, 9]. These studies suggest that glucose metabolism is involved in determining the phenotype of macrophages and thereby regulating inflammatory responses. However, the underlying mechanism is not fully understood.

There are several candidates implicated in macrophage polarisation in response to changes in glucose levels: 1) ATP-sensitive K^+^ channels [10], 2) flux through the hexosamine biosynthesis (HB) pathway [11], 3) AMP-activated protein kinase [12] and 4) activated PKC [13]. Previously, we have shown that exposure of macrophage cell lines RAW264.7 and J774.2 to high levels of glucose increases the β-catenin protein level via the HB pathway [14]. β-catenin is a major component of adherens junction, which connects cadherin to α-catenin protein. Wnt/β-catenin pathway is known to promote M2 macrophage polarization leading to kidney fibrosis [15] and IL-4 induced multinucleated giant cell formation [16]. β-catenin also stabilizes α-catenin by preventing the proteasomal degradation [17]. Our previous studies found that glucose stimulation increases the α-E catenin level in rat pancreatic β-cell models (INS-1E and INS-832/3) [18].

In the current study, we sought to determine whether changes in glucose levels regulate α-E catenin levels in RAW 264.7 macrophages and to assess impacts this could have. We found that changes in glucose levels in RAW 264.7 rapidly alter the level of α-E catenin via a mechanism that requires activity of the HB pathway. We show changes in α-E catenin are associated with changes in expression of pro-inflammatory marker genes. Taken together, these data suggest that α-E catenin protein is regulated by the HB pathway and this contributes to changes in M1/M2 macrophage polarisation mediated by changes in glucose levels.

## Material and methods

### Cell culture

RAW264.7 cells were maintained in RPMI 1640 medium supplemented with 10% (v/v) newborn calf serum, 100 units/ml penicillin and 100 μg/ml streptomycin (All from Life Technologies). Cells were cultured on 12 well format and used for experiment when they are >90% confluent. Cells were serum starved overnight in RPMI media in the presence of 0.5 mM glucose and treated with glucose, glucosamine or other inhibitors as indicated. All the reagents were purchased from Sigma-Aldrich.

### Cell lysate preparation

After treatments, cells were rinsed twice with 0.5 ml of ice-cold 1xPBS and lysates were collected in buffer containing 20 mM Tris–HCl (pH 7.5), 150 mM NaCl, 1 mM EDTA, 1 mM EGTA, 1% Triton X-100, 2.5 mM sodium pyrophosphate, 1 mM β-glycerol phosphate, 1 mM vanadate, 100 mM NaF, 1 mM 4-(2-aminoethyl) benzenesulfonyl fluoride hydrochloride (AESBF), 4 µg/ml leupeptin, and 30 µM N-[N-(N-Acetyl-L-leucyl)-L-leucyl]-L-norleucine (ALLN), 4 µg/ml aprotinin, 0.4 µg/ml pepstatin. Cell lysates were centrifuged at 16 100×***g*** for 10 min, and supernatants were subjected to polyacrylamide gel electrophoresis for Western blotting.

### Western blot analysis

After protein transfer, nitrocellulose membranes were incubated with primary antibodies against α-E catenin (1: 1000; Cell Signalling Technologies, catalogue number # 240) and α-tubulin (1: 20 000; Sigma–Aldrich, catalogue number T6074) at 4°C. After overnight incubation with primary antibodies, membranes were washed and incubated with respective secondary antibodies anti-rabbit IgG HRP (1: 10 000 Santa Cruz biotechnology) or anti-mouse IgG HRP (1: 20 000; Sigma–Aldrich) for 1 hour at room temperature and developed with Clarity™ Western ECL substrate (Bio-Rad Laboratories).

### siRNA transfection

α-E catanin siRNA was transfected to RAW 264.7 macrophages using Nepagene electroporator as per instruction manual. Briefly 250 nM of siRNA was mixed with 1 x10 ^7^ cells/100 μl of RAW 264.7 cells in Opti-Mem and electroporated using below conditions. (Pouring pulse conditions are voltage at 175 V, pulse length 5 ms,, pulse interval 50 ms, 2 pulses, 10% decay rate and + polarity. Transfer pulse conditions are 20V voltage, 20 ms pulse length, 50 ms pulse interval, 5 pulse, 40% decay rate. Cells were used for experiments 48-72 hours after transfection.

### LPS stimulation

72 hours after siRNA transfection, RAW 264.7 cells were treated with 100 ng/ml LPS for 8 hours in the presence of either 0.5 mM Glucose or 20 mM Glucose in serum free media. After treatments cells were washed with 1xPBS and used for RNA isolation.

### Real-Tim PCR (qPCR)

RNeasy Mini Kit (Qiagen) was used to extract total RNA from cells. Samples were then treated with DNase I (Life Technologies) to eliminate genomic DNA. cDNA synthesis (reverse transcription) was performed by using the qScript cDNA Super Mix kit (Dnature) with the same amount of RNA added for all samples. The cDNA synthesis conditions were as follows: 5 min at 25°C > 30 min at 42°C > 5 min at 85°C > hold at 4°C. cDNA samples were loaded onto a 394 well PCR plate prior to qPCR. In addition, DNase-free water was included as non-template control. Each qPCR reaction consisted of 5 μL qScript cDNA Super Mix, 5 ng cDNA (equivalent to RNA added) and 400 nM primers and water (up to 10 μL). PowerUp SYBR Green Master Mix (Applied Biosystems) was used for qPCR reactions. The qPCR conditions were as follows: 2 min at 50°C > 10 min at 95°C > 40 cycles of (15 sec at 95°C, 1 min at 60°C) > 15 sec at 95°C > 1 min at 60°C > 15 sec at 95°C. The double delta Ct analysis was performed to calculate the gene expression fold change in relation to the control. A single distinct peak in melt curve was checked to validate the qPCR reaction in each well.

### Statistical analysis

Results are presented as means ± S.E.M. with the number of experiments indicated in the legend. Statistical significance was assessed using student’s t-test or two-way ANOVA as indicated in figure legends. Statistical significance is displayed as * *P* < 0.05 or ** *P* < 0.01. Statistical analyses were performed using statistical software package GraphPad Prism 6.0 (GraphPad Software Inc.)

## Results

We found that high levels of glucose cause an increase of α-E catenin protein levels in the RAW 264.7 macrophage model (Figure 1 A-B), consistent with our previous findings in rat pancreatic β-cells [18]. As we have previously found that β-catenin levels increased with glucose stimulation in macrophage cell lines through the HB pathway [14], we speculated that the same pathway might account for the glucose-induced upregulation of α-E catenin. To investigate this possibility, RAW 264.7 cells were treated with glucosamine. Glucosamine is converted to glucosamine-6-phosphate (GlucN-6-P), bypassing the first three steps required for glucose in the HB pathway (Figure 1 C). Glucosamine increased α-E catenin levels in macrophages in a similar manner to that seen with high glucose (Figure 2A-B). The conversion of fructose-6-phosphate to GlucN-6-P (the rate limiting step of the HB pathway) is regulated by glutamine: fructose-6-phosphate-amidotransferase (GFAT) enzyme, which requires glutamine as a co-substrate [19]. Therefore, to further confirm that the HB pathway is responsible for modulating the level of α-E catenin, we treated RAW 264.7 cells with high glucose in the absence of glutamine and observed no significant change in α-E catenin level (Figure 2C-D). In line with this finding, when GFAT enzyme was inhibited with azaserine, the glucose effect on α-E catenin levels were attenuated (Figure 3A-B). The HB pathway mediates the glycosylation of proteins, which is important for protein stability, protein activity, cell-cell communication and signal transduction [20, 21]. Next, we treated cells with tunicamycin, which is an inhibitor of N-linked glycosylation. The glucose effect on α-E catenin was attenuated in the presence of tunicamycin (Figure 3C-D), indicating that the regulation of α-E catenin level relies on N-linked glycosylation. These findings suggest that α-E catenin level in RAW264.7 is regulated via the HB pathway involving N-linked glycosylation.

**Figure 1.**
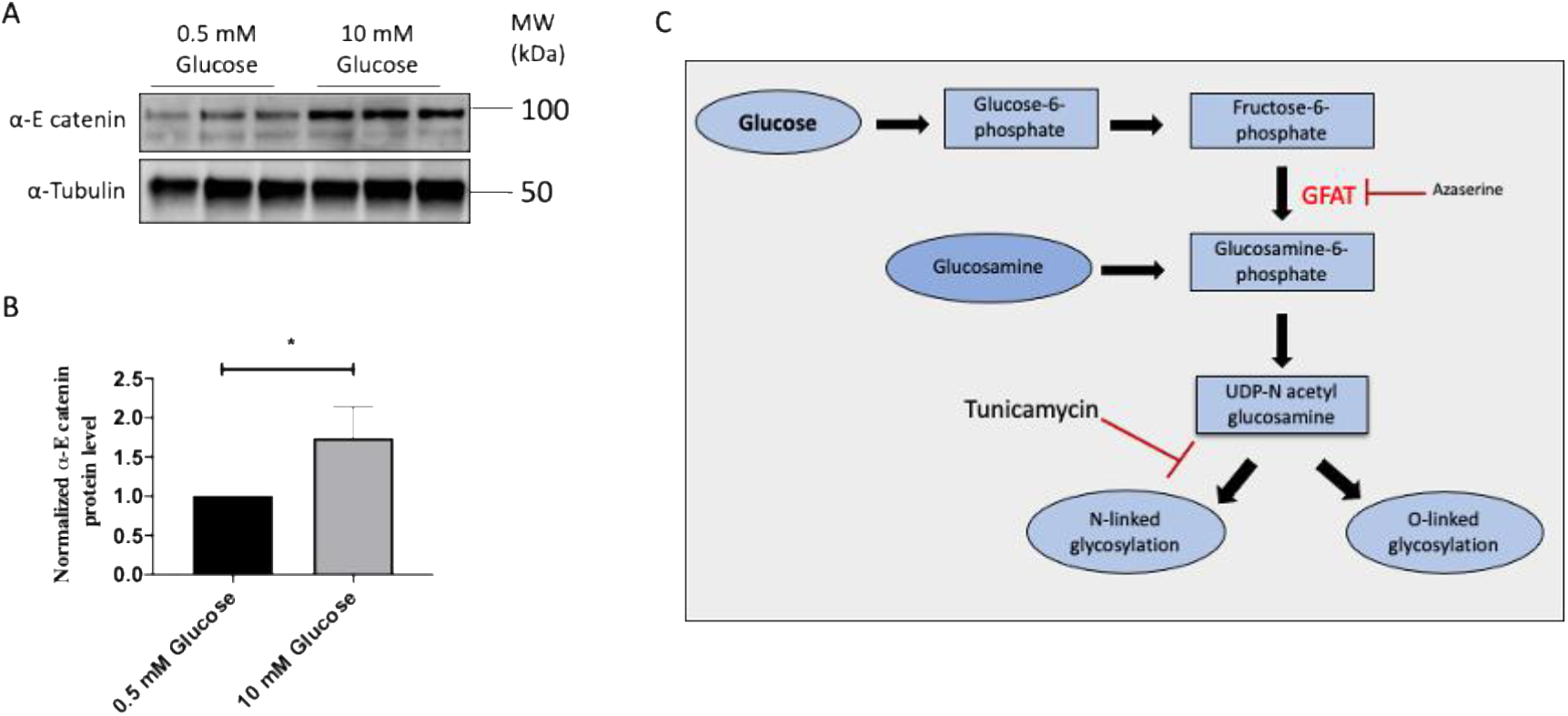
High glucose level increases the α-E catenin level in RAW 264.7 cells. After overnight serum starvation in RPMI media with 0.5 mM glucose, RAW 264.7 cells were treated with either 0.5 mM or 10 mM glucose for 2 hours. (A) Cell lysates were subjected to the Western blot analysis using α-E catenin and α-tubulin antibodies. (B) Densitometry analysis showing protein expression of α-E catenin normalised to α-Tubulin. Data represent mean ± S.E.M of three independent experiments. The unpaired t-test was used to assess statistical significance *P <0.05.

**Figure 2.**
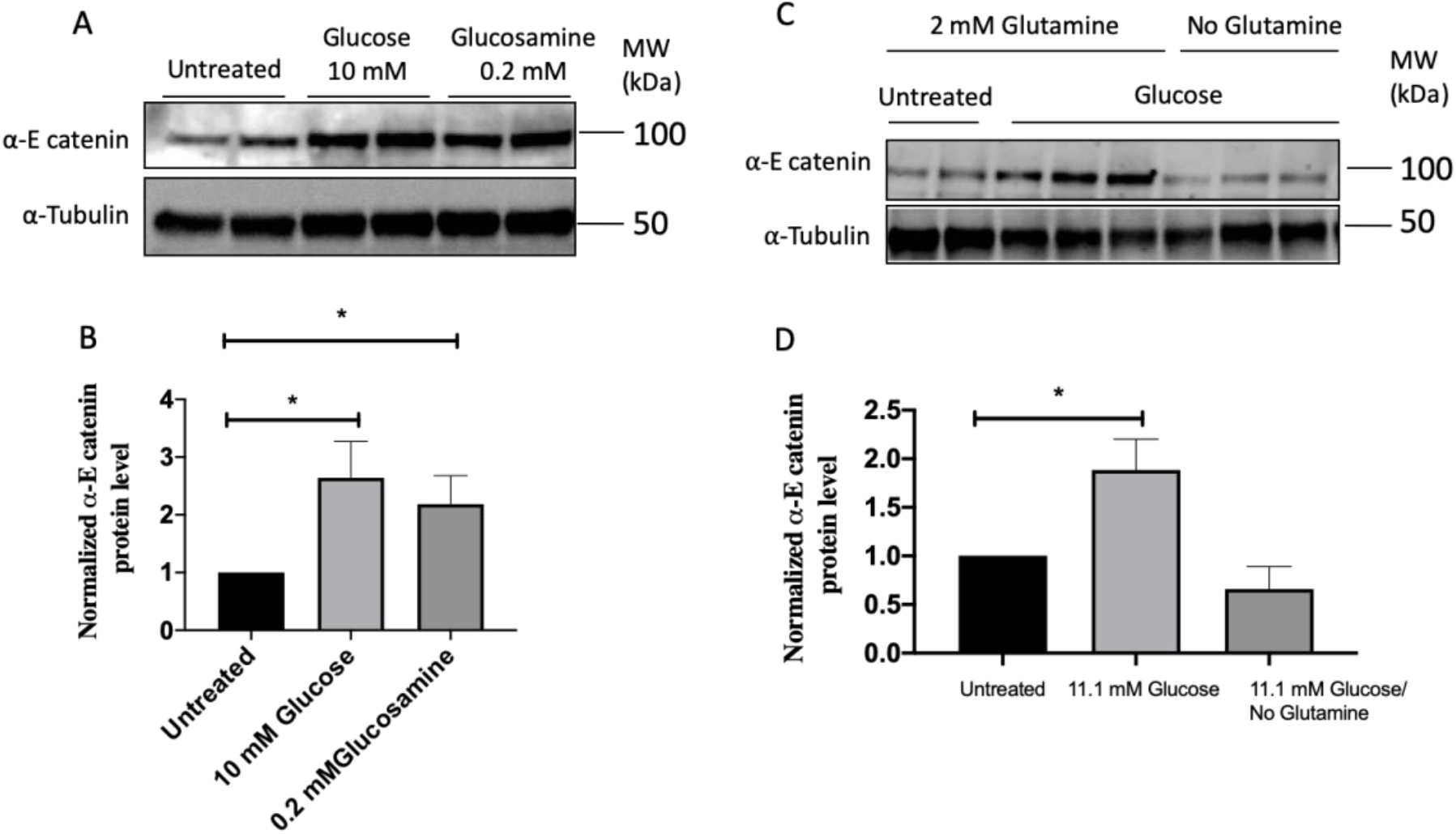
Glucosamine increases the α-E catenin level in RAW 264.7 cells and Glutamine is required for glucose induced increase of α-E catenin. RAW 264.7 cells were treated with indicated concentrations of glucose or glucosamine for 2 hours after overnight serum starvation in the presence of 0.5 mM glucose (A) Cell lysates were subjected to the Western blot analysis using α-E catenin and α-Tubulin antibodies. (B) Densitometry analysis of α-E catenin protein expression in western blot image relative to the α-Tubulin. RAW 264.7 cells were treated with indicated concentrations of glucose in the presence or absence of glutamine for 2 hours. (C) Cell lysates were subjected to the Western blot analysis using α-E catenin and α-Tubulin antibodies. (D) Densitometry analysis showing protein expression of α-E catenin normalised to α-Tubulin. Data represent mean ± S.E.M of at least four independent experiments. The unpaired t-test was used to assess statistical significance *P <0.05

**Figure 3.**
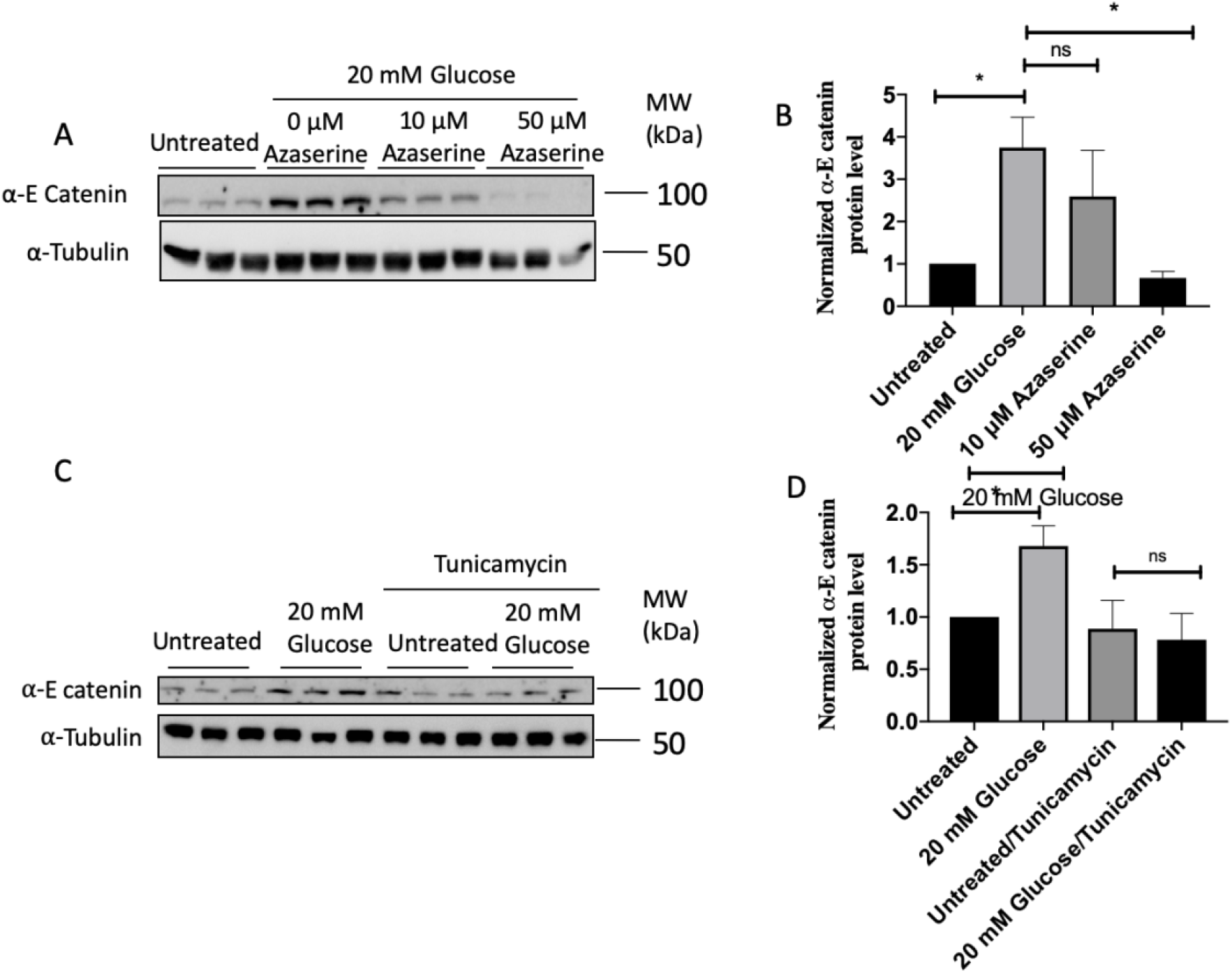
Azaserine and Tunicamycin attenuate glucose induced increase of α-E catenin. After overnight serum starvation in RPMI media with 0.5 mM glucose, RAW 264.7 cells were treated with 20 mM glucose in the presence or absence of azaserine for 4 hours. (A) Cell lysates were subjected to the Western blot analysis using α-E catenin and α-Tubulin antibodies. (B) Densitometry analysis showing protein expression of α-E catenin normalised to α-Tubulin. Data represent mean ± S.E.M of three independent experiments. The unpaired t-test was used to assess statistical significance *P <0.05 RAW 264.7 cells were treated with 20 mM glucose in the presence or absence of tunicamycin (10 μg/ml) for 2 hours. (C) Cell lysates were subjected to the Western blot analysis using α-E catenin and α-Tubulin antibodies. (D) Densitometry analysis showing protein expression of α-E catenin normalised to α-Tubulin. Data represent mean ± S.E.M of four independent experiments. The unpaired t-test was used to assess statistical significance *P <0.05

We next investigated the role of α-E catenin in the function of the RAW cells using siRNA to reduce α-E catenin. We electroporated RAW 264.7 cells with an α-E catenin siRNA, which led to at least 50% reduction in levels of α-E catenin (Figure 4 A). We then treated cells with either low (0.5 mM glucose) or high glucose (20 mM glucose) in the presence or absence of LPS. We found that decreased expression of *Ctnna1* attenuates glucose-induced upregulation of *IL-1β* under the LPS stimulated condition (Figure 4 B). In addition, we observed the decreased expression of *IL-12* in the α-E catenin knockdown samples, compared to the control, when treated with low glucose under the LPS stimulated condition (Figure 4C). Interestingly, we observed a significant decrease in *IL-12* upon high glucose challenge in the control samples treated with LPS, while this did not occur in the knockdown samples (Figure 4 C). Under LPS stimulated conditions, we did not observe any effect of α-E catenin knockdown on *TNF-α* at either low or high glucose conditions (data not shown).

**Figure 4.**
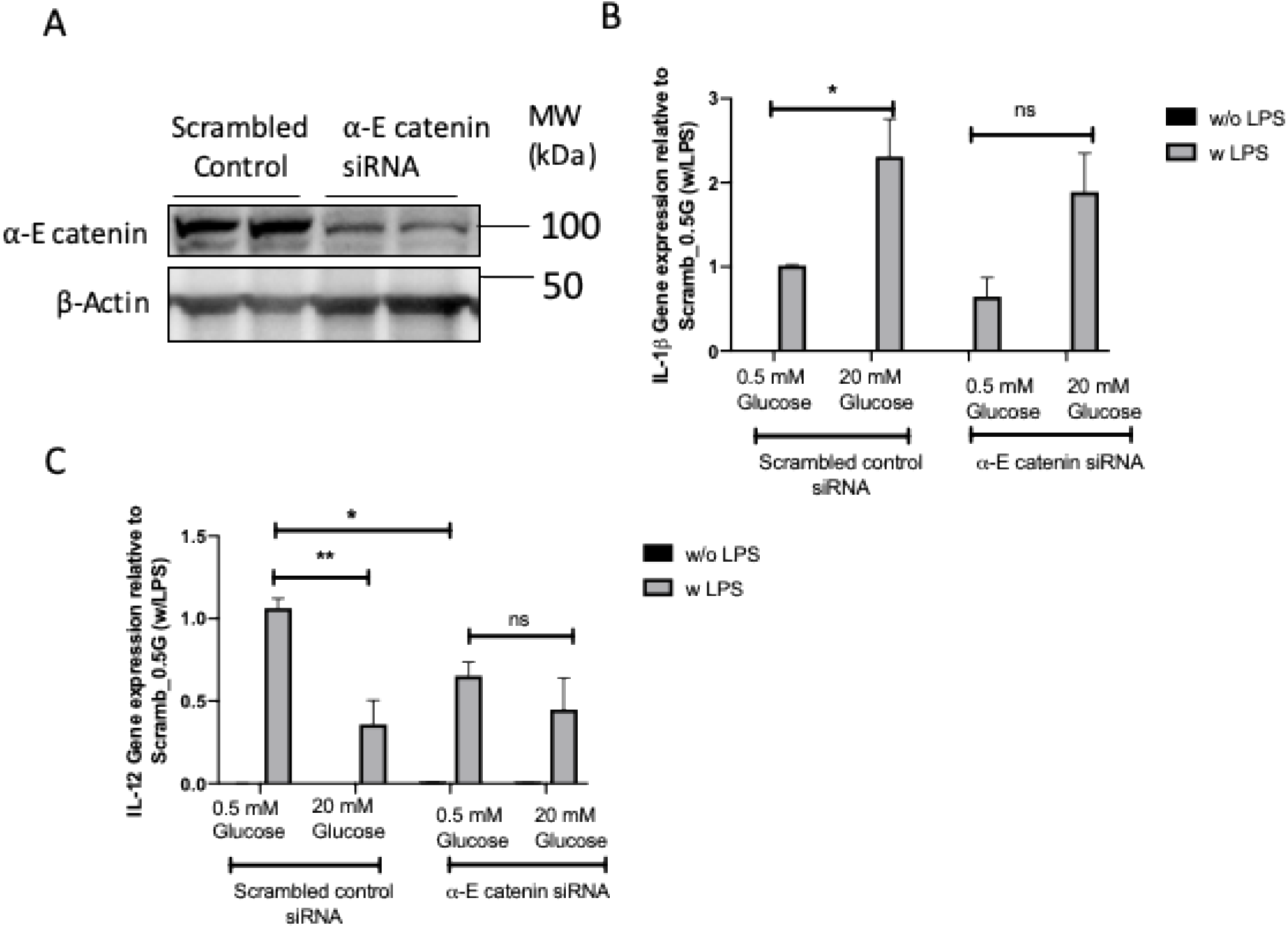
α-E catenin knockdown attenuates glucose induced increase of *IL-1β* and basal level of *IL-12*. 48 hours after siRNA transfection, RAW264.7 cells were either untreated or treated with LPS in the presence of 0.5 mM or 20 mM glucose for 8 hours. (A) Cell lysates were subjected to the Western blot analysis using α-E catenin and α-Tubulin antibodies (B) qRT-PCR showing relative gene expression of *IL-1β* and *IL-12*. Data represent mean ± S.E.M of three independent experiments, assessed by two-way Anova. *P <0.05, ** P <0.01

## Discussion

Macrophages have phenotypic plasticity which is an important for their function in inflammation. High glucose levels favour macrophage polarization towards the M1 phenotype, characterised by the secretion of pro-inflammatory cytokines which induces inflammation of surrounding environment. The most novel finding of the current study is identification of a previously unreported role of α-E catenin in macrophages by showing pro-inflammatory genes *IL-1β* and *IL-12* being regulated by α-E catenin levels. This implicates α-E catenin as a new component of mechanisms involved in switching macrophages between M1 and M2 states. We find that that α-E catenin protein level is increased upon glucose stimulation using the RAW 264.7 cell line as a macrophage cell model via the HB pathway which could thus contribute to the changes in macrophage function known to be brought on by changes in glucose levels [1, 2].

One outcome of the HB pathway is protein glycosylation and a lack of proper protein glycosylation results in accumulation of misfolded proteins in the endoplasmic reticulum (ER) which results in ER stress [22]. Thus, the finding that the glycosylation inhibitor tunicamycin blocks glucose induced increases in α-E catenin in our study suggests normal ER function would be required for this mechanism of α-E catenin regulation. It is notable that a separate proteomic approach has also shown that ER stress also reduces both α-catenin and β-catenin protein levels in HeLa cells and that ER stress was reducing catenin levels by increasing rates of proteasomal degradation [22]. Proteasomal turnover of α and β-catenin is associated with increased CK1 and GSK3 mediated phosphorylation of β-catenin at ser37/thr41 [23, 24]. Tunicamycin regulates GSK3 in the proteomic study [22] and studies in other cell types also show that tunicamycin activates GSK3 which would be consistent with reductions in α and β-catenin [25-28]. However, we don’t see any changes on GSK3β levels or phosphorylation status despite seeing a reduction in β-catenin Ser37/Thr41 phosphorylation [14]. CK1 activity has also been reported to a change in response to changes in glucose levels in some c ell types [29] and tunicamycin may increase CK1 activity [30] although it is not known if this occurs in macrophages. Therefore, the mechanisms linking changes in glucose levels to changes in α-E catenin levels remain to be elucidated.

The functional implications of our findings remain to be determined but could have implications for pathology associated with type-2 diabetes where glucose levels are constantly elevated [2]. In our previous work, we also have demonstrated that α-catenin inhibits insulin secretion in pancreatic β-cells [18]. Type-2 diabetes is characterized by β-cell failure [31] in which chronic inflammation involving macrophages has been implicated [32, 33]. Thus, it will be worth investigating how glucose induced changes in α-E catenin in both β-cells and macrophages might combine in promoting progression of this disease.

## Declarations

### Availability of data and material

No large data sets associated with this study. All raw data for western blots and qPCR available upon request.

### Code availability

Not applicable

### Author Contribution

Waruni Dissanayake, Brie Sorrenson, and Peter Shepherd contributed to the experimental design. Waruni Dissanayake and Jin Kyo Oh performed the experiments. Waruni Dissanayake, Jin Kyo Oh and Peter Shepherd analysed and interpreted the data. Waruni Dissanayake and Peter Shepherd wrote the paper.

### Ethics approval

Not applicable

### Consent to participate

Not applicable

### Consent for publication

Not applicable

